# Lactic Acid Production by *Clostridium acetobutylicum* and *Clostridium beijerinckii* Under Anaerobic Conditions Using a Complex Substrate

**DOI:** 10.1101/2021.02.23.432258

**Authors:** Mattiello-Franisco L., Vieira F.M., Peixoto G., Mockaitis G.

## Abstract

High societies consumption, elevated residues generation and environmental awareness strengthen alternatives solutions for bioprocess’ residues. This study investigated the production of volatile acids from a complex substrate, which intends to be replaceable in the future by vinasse of sugar cane, in anaerobic reactors operated in triplicates at 35 °C. Two different inoculum were studied: *Clostridium acetobutylicum* ATCC 824 and *Clostridium beijerinckii* ATCC 25752. The nutrient medium had as carbon source a complex substrate containing sucrose without addition of vitamins, buffer solution and micronutrients. The experiment was conducted in the variation of F/M-ratio (food-to-microorganisms) by increasing substrate concentration.

The concentration of sucrose in the complex substrate were 5.2 g·L^-1^ (conditions of 10,000 mg O_2_· L^-1^ in terms of COD) and 10.5 g· L^-1^ (conditions of 20,000 mg O_2_·L^-1^ in terms of COD), keeping the initial concentration of inoculum in 500 mg SVT·L^-1^. Cultures *C. acetobutylicum* and *C. beijerinckii* resulted in high lactic acid production. Concentrations of COD of 10,000 mg O_2_· L^-1^ produced optimum lactic acid of 3,331 mg·L^-1^ and 5,709 mg·L^-1^ with respectively C. *acetobutylicum* and *C. beijerinckii.* Moreover, cultures C. *acetobutylicum* and *C. beijerinckii* with 20,000 mg O_2_· L^-1^ concentrations in terms of COD produced optimum lactic acid of 6,417 mg· L^-1^ and 7.136 mg·L^-1^ respectively. There was repeatability in the reactors when considering level of significance of 0.05, independent of the concentration and inoculum used.

## 1 INTRODUCTION

The replacement of fossil fuels by other sources of renewable energy grows worldwide, mainly because of the pressure of policies to preserve the environment. Processes that use biomass as raw material rather than non-renewable source are called biorefinery. An example of biorefinery in Brazil is sugar cane mills. The apparent antagonism between the environmental issue and the increased productivity can affect the economic viability of some industrial processes. Thus, an expanded concept of biorefinery aimed at a sustainable process is the use of all material flows present in a biological process. In sugarcane mills, the new concept of biorefinery as a sustainable process is established when using its residues, such as vinasse, for the generation of products with higher added value, such as acids and solvents.

Ethanol from sugarcane makes up a significant part of Brazil’s bioenergy matrix. In the 2015, it was showed that 43.5% of the Brazilian energy matrix comes from renewable energy, 18.1% of which is sugarcane [1]. In the 2014/2015 crop, Brazil produced 634.8 million tons of sugarcane in 9 million hectares. Specifically, for ethanol production were 361 million tons of sugarcane, equivalent to 56.9% of total production [2]. The State of São Paulo has a strong influence in this sector, representing 49.4% of the country’s ethanol production. In the 2016/2017 harvest, the State of São Paulo produced approximately 365.9 million tons of sugarcane [3]. Even though ethanol is an interesting alternative as biofuel, it continues to be a topic of improvements in the production process. At the Brazilian mills, on average, one ton of sugar cane produces 280 kg of bagasse [4] and 800 to 1,000 liters of vinasse [5]. Sugarcane vinasse is the liquid residue generated on a high scale in the ethanol production process. It is reported that, on average, in order to produce 1 liter of ethanol 10 – 15 liters of vinasse are generated which may vary depending of the material and technology used [6]. Since vinasse is a largely generated byproduct of ethanol production it is an example of a range of biomass which have been studied as an alternative substrate in anaerobic bioprocessing to produce added value products. It is constituted of organic solids and minerals, which may provide a rich culture medium for bioprocesses [7].

Due to the diversity of the composition during the harvests, some researchers choose to use a synthetic solution with characteristics similar to the biomass for conducting research. A synthetic effluent [8] with characteristics similar to the soluble fraction (1: 5) of sugarcane vinasse was used to evaluate the efficiency of COD and sulphate removal in the anaerobic fixed-bed and downflow (DFSBR) together with an acid solution rich in iron ore [8]. The substrate composition was defined based on the characterization of the real vinasse.

Anaerobic digestion is a process that occurs in the absence of oxygen and transforms various forms of complex organic matter (carbohydrates, proteins and lipids) into simpler products (such as carbon and methane) by the metabolism of a consortium of different microorganisms (SPEECE, 1996). Among the several metabolic routes present in the anaerobic process, the ABE (acetone, butanol and ethanol) fermentation was discovered in the 1920s by Chaim Weizmann. Effective anaerobic bacteria in the ABE fermentation process are those of the genus *Clostridium*, belonging to at least 60 species of bacteria of this genus. The anaerobic bacterium *C. acetobutylicum* produces naturally acetone, butanol and ethanol in a ratio of 3: 6: 1 from the pure glucose substrate [9]. It is emphasized that the metabolic sequence of *C. acetobutylicum* in the ABE fermentation has two phases: acidogenesis and solventogenesis [10]. In the acidogenic process cell growth occurs and the production of organic acids (acetic acid and butyric acid) with the consequent decrease of the pH of the medium to 4.5. Then, in the solventogenic process, the acids produced in the previous phase are in high concentrations allowing them to be processed in solvents. Although ethanol production is also part of the solventogenic process, it is considered that this route can occur independently of the production of acetone and butanol [11].

Even though, the metabolic pathway of pyruvate to lactate shows as a deviation from ABE fermentation, lactate formation is a route of paramount importance and widely studied. The enzyme D-lactate dehydrogenase (LDH) is present in the reaction of pyruvate to lactate when using *C. acetobutylicum* ATCC 824. The enzyme D-Lactate dehydrogenase (LDH) has the cofactor fructose-1-6-diphosphate, and therefore is an enzyme with allosteric activation [13, 14]. The anaerobic digestion of organic compounds to lactate instead of methane has some advantages due to the lower hydraulic retention time (HRT) which it is reported as less than 5 days against 28 days in single stage reactors [14]. In addition, lactic acid has applications in pharmaceutical and chemical industries, besides food and beverage sector, and it can be transformed in the biodegradable polymer poly(lactic acid) (PLA) [15].

This study is an experimental investigation aiming the production of acids and solvents from a complex substrate (with characteristics based on sugarcane vinasse) and the two different inoculums: *C. acetobutylicum* and *C. beijerinckii*.

## 2 MATERIAL AND METHODS

### 2.1 Experimental procedure

The experimental was done in batch reactors maintained under constant agitation in shakers (100 min^-1^) at 35°C during 720 hours. Those batch reactors were Duran^®^ flasks (500 mL) with reactional volume of 250 mL and initial pH of 6.0. N_2_-gas was fluxioned 3 times of 8 min each to keep an anaerobic environment. The batch reactors were structured into two inoculums (*C. acetobutylicum* and *C. beijerinckii*) with substrate concentration of 10,000 and 20,000 mg O_2_·L^-1^ in terms of chemical oxygen demand (COD). The main parametron used was the ratio F/M-ratio, maintaining constantly the inoculum concentration of 500 mg SVT·L^-1^. This ratio of inoculum was determined according to the variation of 2 to 4% used in an industrial scale [11]. Therefore, concentration of 10,000 mg O_2_·L^-1^ in terms of COD refers to 5% and concentration of 20,000 mg O_2_·L^-1^ in terms of COD refers to 2,5%. The experimental design of the research is detailed in Table 1.

**Table 1.** Experimental design of the influence of complex substrate concentration variation on each inoculum

| Organic matter<br>(mg O<sub>2</sub>·L<sup>-1</sup>) | F/M<br>(mg O<sub>2</sub>·mg SVT<sup>-1</sup>) | *C. acetobutylicum* | *C. beijerinckii* |
| --- | --- | --- | --- |
| 10,000 | 20 | Ac-10 | Bj-10 |
| 20,000 | 40 | Ac-20 | Bj-20 |

As shown in Table 1, the inoculums *C. acetobutylicum* and *C. beijerinckii* will be referred respectively to the abbreviation “Ac” and “Bj” in order to simplify the discussion of the results. In each condition the reactor cycle started after inoculation and the end of the test occurred in 720 hours. All assays of the experimental conditions previously described were conducted in triplicates.

### 2.2 Composition of substrate

The composition of the substrate used in this research was based on the macro and micronutrients employed by Godoi et al. (2017) [8].

The composition of substrate used in these experiments aims to approximate the complex substrate to the sugar cane vinasse. The study of the metabolism of a synthetic substrate rather than the actual substrate aims at the future application of this fundamental study in the biological processing of this residue. In this sense, it is important that the substrate used in this investigation is not a variable source of concentration and composition, which would impair the conclusions obtained. Table 2 shows the average concentrations of sucrose (carbon source) with their respective standard deviation used in the tests of 10,000 mg O_2_·L^-1^ and 20,000 mg O_2_·L^-1^ in terms of COD.

**Table 2.**
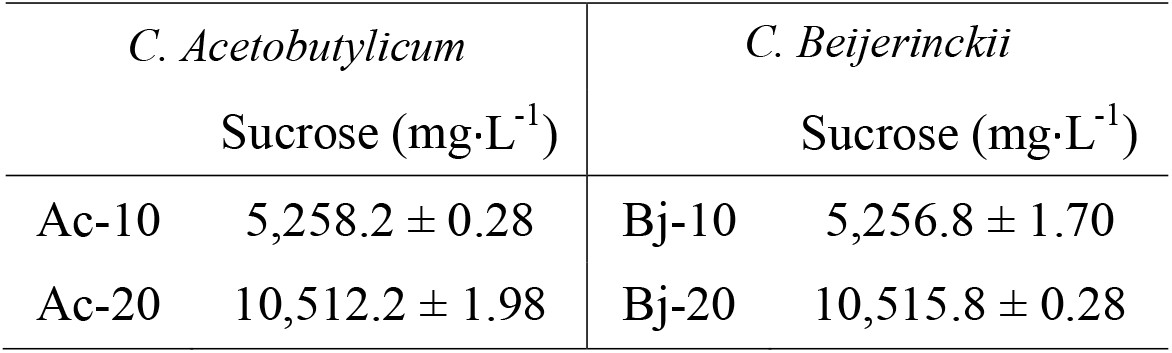
Concentration of sucrose (mg·L^-1^) in conditions of 10,000 mg O_2_·L^-1^ and 20,000 mg O_2_·L^-1^ in terms of COD with *C. acetobutylicum* and *C. beijerinckii.*

It can be seen from the standard deviation shown in Table 2 that the triplicates had little variation in relation to the sucrose concentration, which allows comparisons of results in terms of metabolites production.

### 2.3 Inoculum

The pure cultures of *C. acetobutylicum* ATCC 824 and *C. beijerinckii* ATCC 25752 were obtained through a microorganism collection of the André Tosello Foundation - Campinas, SP. Both strains were stored in ampoules in lyophilized form. The strains were reactivated and the contents of the ampoules were transferred to a test tube containing 5 ml of liquid culture medium (RCM - Reinforced Clostridium Medium). Multiplication of each culture was performed in a 2L Erlenmeyer flask with 1L RCM medium kept in an oven at 35 ° C for at least 7 days until the turbidity of the medium stabilized. Monitoring of the growth curve of the cultures was performed by analysis of turbidity (λ = 500 nm) and total volatile solids.

### 2.4 Monitoring analysis

The pH and organic matter concentration determined according to Standard Methods [16]. The following variables were also monitored: carbohydrates [17], organic acids (Penteado et al., 2012) and alcohols (Penteado et al., 2012 and Pavini, 2017).

The liquid phase composition analyzes were determined by a modular Shimadzu^®^ liquid chromatograph (HPLC), using an LC-10AD pump system, a CTO-20A column oven, a SCL-10A controller, a luminous matrix detector (PDA - Photo Diode Array), adjusted for scanning comprising a wavelength range of 190 to 370 nm (UV region), with 1 nm step, the chromatogram being read at 210 nm and a RID-10A refractometer detector with cell temperature of 35°C. The fixed phase comprised a BIO-RAD Aminex^®^ HPX-87H 3000 x 7.8 mm column, with pre-column of the same type, operating at a constant temperature of 43°C. The eluent used was a 0.005 mol·L^-1^ H_2_SO_4_ solution at the flow rate of 0.5 ml·min^-1^ and the volume injected was 100 μL. The integration and identification of peaks was performed using Shimadzu Class-VP^^®^^ software version 5.032.

Analysis for butanol determination were carried out in the CEMPEQC laboratory using Thermo Electron Corporation (Thermo Scientific) model Trace GC Ultra, coupled with a TriPlus AS (Thermo Scientific) automatic sampling system and SGE Analytical Science liquid syringe with 10 μL capacity. The column used was ZB-WAX (30 m x 0.25 mm x 0.25 μm, Phenomenex), with split/splitless injector, flame ionization detector (FID) and helium drag gas (He). The extraction of the analytics from the aqueous sample was performed according to the Pavini method (2017) by adding 0.555 g of sodium sulfate in 2.0 ml Eppendorf with two fractions of 600 μL of sample and 400 μL of solution of 1-octanol containing Internal Standard at the concentration of 1,000 mg·L^-1^. After homogenization of the Eppendorf content for 30 seconds on vortex tubes, 900 μL of the supernatant (organic fraction) was removed for analysis on the gas chromatograph.

### 2.5 Kinetic analysis

The residual first order kinetic model from the simplification of the kinetic model proposed by Monod was used due to the low substrate concentrations. Equation 1 shows the model used to evaluate this study [18].

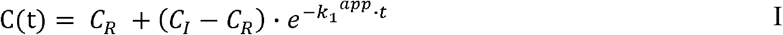

Where C(t) is the concentration of sucrose (mg·L^-1^); C_R_ is residual sucrose concentration (mg·L^-1^); C_I_ is the initial concentration of sucrose (mg·L^-1^); t is the time of the experiment (hours) and k_1_^app^ is the apparent kinetic constant (h^-1^).

For the calculation of the lactic acid production it was used the simple dose-response sigmoid model, presented in Equation 2.

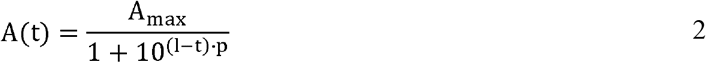

In Eq. 2, A(t) is the accumulated production of the product after a certain time t of fermentation (mg·L^-1^); A_max_ is the maximum production of the product (mg·L^-1^); l is the time to reach the maximum production speed (h); t is the time of the experiment (h) and p is the average rate of production of the product in the exponential growth phase (mg · L^-1^·h^-1^).

The Levenberg-Marquadt interaction algorithm of Microcal Origin^®^ v 8.1 software was used as an adjustment tool for kinetic parameters.

## 3 RESULTS AND DISCUSSION

Acids and solvents production from a complex substrate are influenced by reactors environmental. The fermentation process is dependent on a variety of conditions such as nutrient shortage and pH, the metabolism is highly influenced by these parameters and the fermentation can be favored in order to produce acids or solvents [19].

The parameters used to evaluate this study were consumption of sucrose and pH profile throughout the experiment.

### Substrate consumption profiles

Both inoculums were able to adapt to the complex substrate as well as to the concentrations evaluated. This behavior can be confirmed by the reduction of the organic matter concentration, expressed in terms of COD, observed in Figure 1. The adjustment of the first order residual kinetic model for sucrose concentration has been referenced by the line.

**Figure 1.**
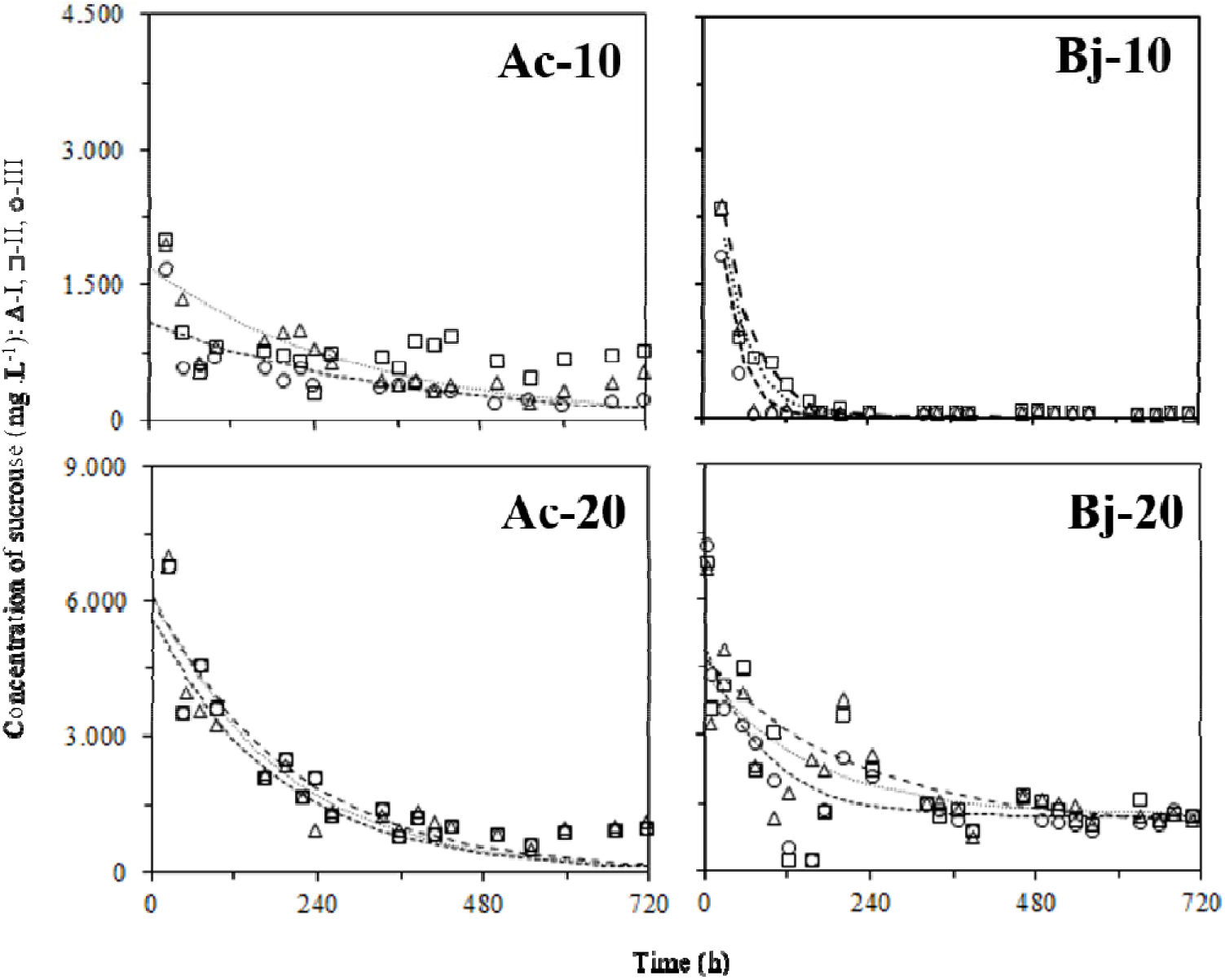
Sucrose concentration profiles with *C. acetobutylicum* and *C. beijerinckii* inoculums at concentrations of 10,000 mg O_2_·L^-1^ and 20,000 mg O_2_·L^-1^.

The first sucrose concentration analysis occurred after 24 hours from experiment start. All the sucrose concentration adjustments presented satisfactory statistical correlation values (R_2_> 0.62), mainly using *C. beijerinckii* (R_2_> 0.85), indicating that the first-order kinetic model was adequat to represent the behavior of the substrate consumption for the both COD concentrations. The value variation of the first order apparent kinetic constants (k_1_^app^) are presented in Table 2 and demonstrated that the increase of the initial complex substrate concentration exerted a favorable effect to the process. Therefore, lower values of the first order apparent kinetic constant (k_1_^app^) were obtained in the replicates of Bj-20 in relation to Bj-10.

| Strain | COD | $k_1^{\text{app}}$ |
| --- | --- | --- |
|  | (mg L<sup>-1</sup>) | (h<sup>-1</sup>) |
| *C. acetobutylicum* | 10,000 |  |
|  | 20,000 |  |
| *C. beijerinckii* | 10,000 |  |
|  | 20,000 |  |

### pH profiles

The pH profiles were performed at COD concentration of 10,000 mg O_2_·L^-1^ and 20,000 mg O_2_·L^-1^ with *C. acetobutylicum* inoculum (conditions Ac-10 and Ac-20) and *C. beijerinckii* (Bj-10 conditions and Bj-20). Each graph on the Figure 2 presents three pH curves for each triplicate (replicate I-Δ, replicate II - □ and replicate III - ○).

**Figure 2.**
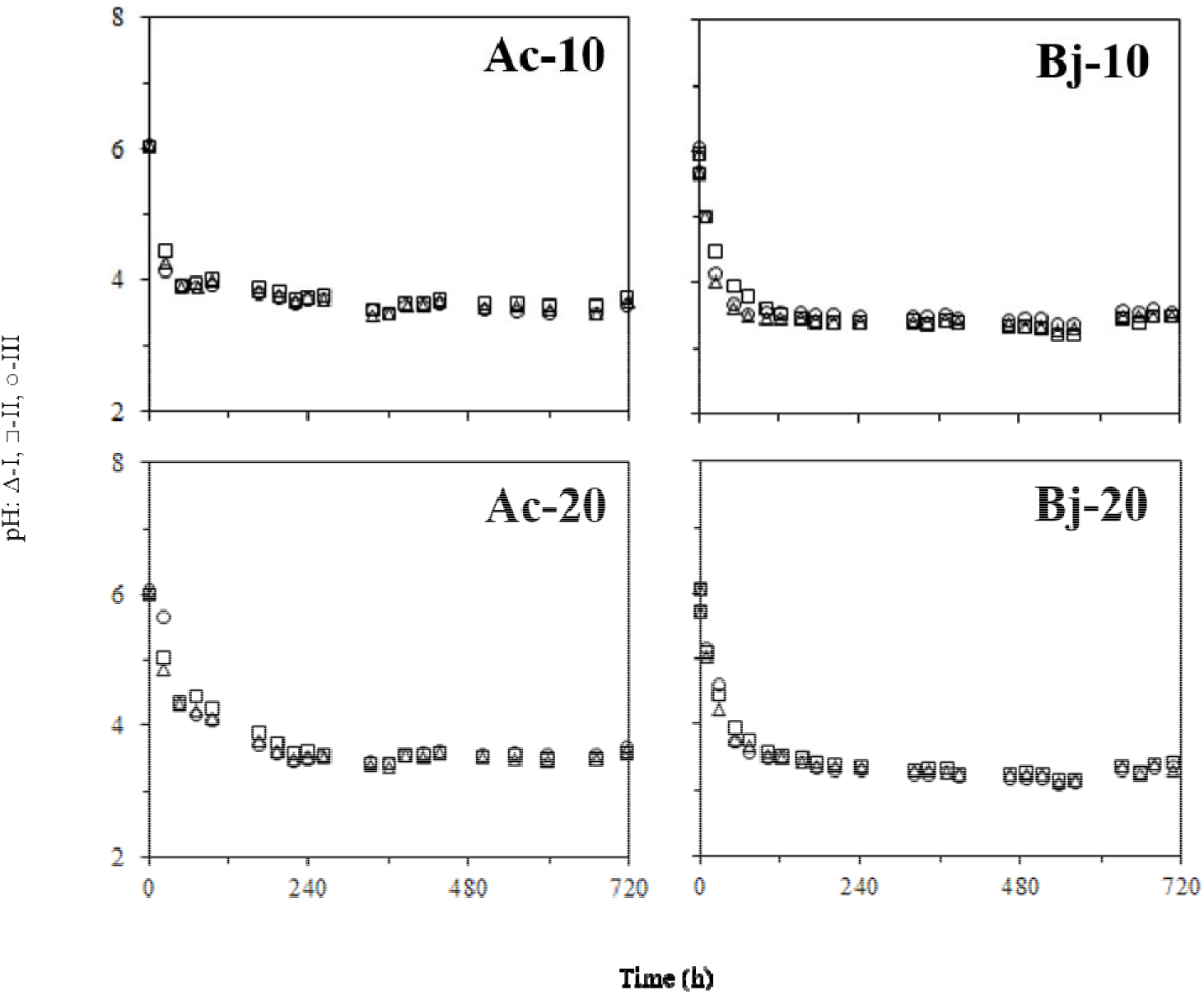
pH profiles in the assays with *C. acetobutylicum* and *C. beijerinckii* inoculums at concentrations of 10,000 mg O_2_·L^-1^ and 20,000 mg O_2_·L^-1^.

It can be observed that the pH declines throughout the experiment, suffering a variation from 6.0, which was set at all batches, to 3.5. Fig. 2 and 3 highlight that the lactic acid production occurred mostly when the pH was declined. The fermentation products were strongly affected by the pH. These observations agree with literature which has already reported that the metabolism of *Clostridium* strains shifts when the pH declines. It was observed that when pH decreased to below 6.0, acetic acid and lactic acid concentration increased, and the acidogenic pathway wa favored. When the pH was reduced to below 5.0, lactic acid was the major fermentation product [20].

**Figure 3.**
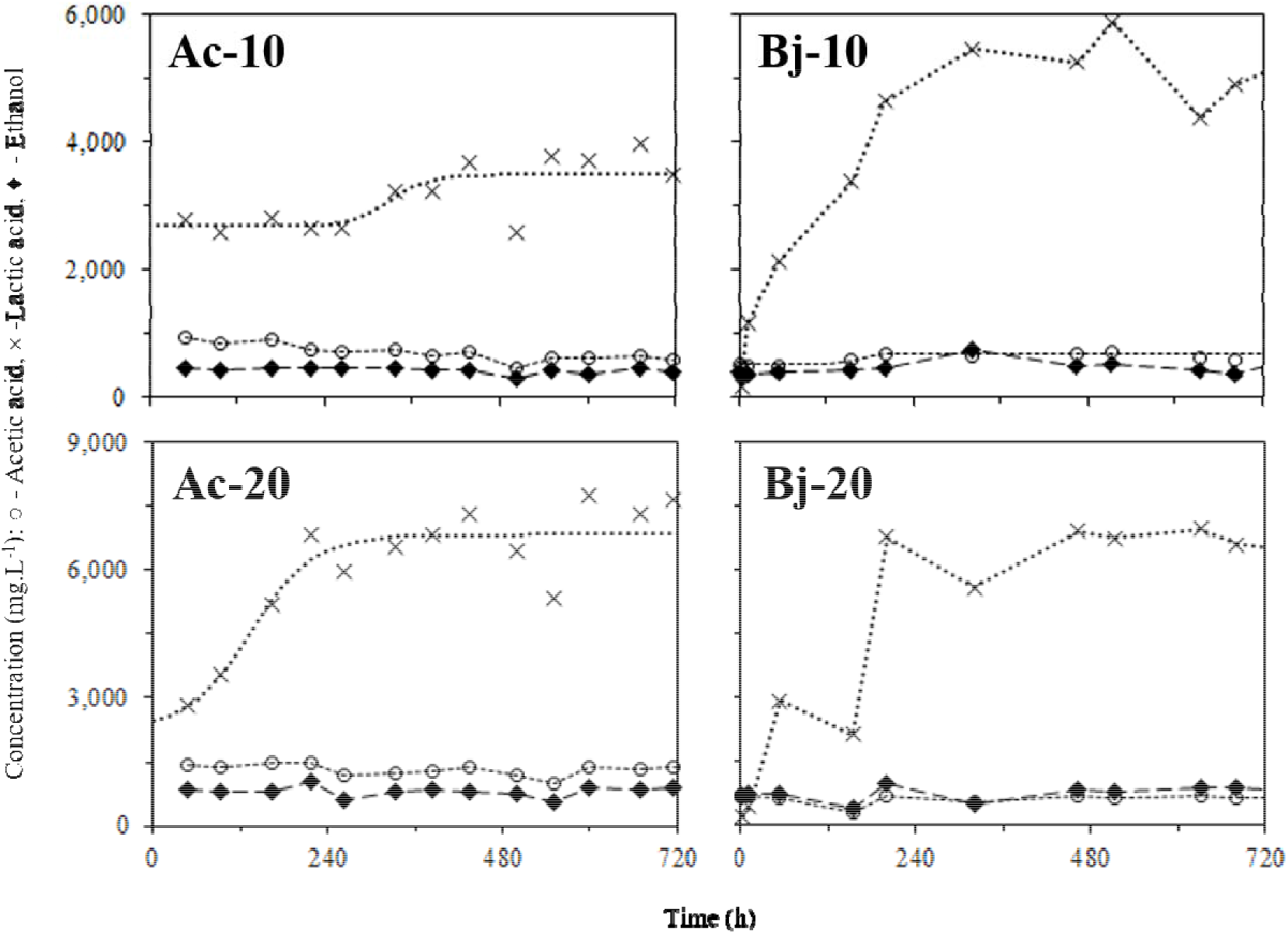
Products from reactors with *C. acetobutylicum* and *C. beijerinckii* inoculums at concentrations of 10,000 mg O_2_·L^-1^ and 20,000 mg O_2_·L^-1^.

### Metabolites production

The metabolites production was detected and monitored throughout the experiment. Figure 3 shows the production of acetic acid (○), lactic acid (×) and ethanol (♦) occurred in one replicate for each condition.

It is observed in Figure 3 that the concentration of ethanol and acetic acid remained stable during the experiment, while lactic acid production was favored. This behavior of the batches can occur due to the decrease of the pH, as already discussed. The carbon source concentration also plays a major role in the experiment performance and can be observed in Fig. 3, since the conditions with higher COD presented higher lactic acid production.

According to Figure 3, a simple sigmoidal curve was performed for lactic acid production using *C. acetobutylicum.* Kinetic parameters for those curves are presented in Table 3. The best adjustment occurred with *C. acetobutylicum* at concentrations of COD of 20,000 mg O_2_·L^-1^.

**Table 3.** Kinetic parameters (*A_max_*, *A_min_*, *l* e *p*) and correlation coefficient (R^2^) related to lactic acid from reactors with *C. acetobutylicum* at concentrations of 10,000 mg O_2_·L^-1^ and 20,000 mg O_2_·L^-1^

| | | Kinetic Parameters | | | | $R^2$ |
| --- | --- | --- | --- | --- | --- | --- |
| | | $A_{max}$<br>(mg·L<sup>-1</sup>) | $A_{min}$<br>(mg·L<sup>-1</sup>) | $l$<br>(h) | $p$<br>(g·L<sup>-1</sup>·h<sup>-1</sup>) | |
| Lactic acid | Ac-10 | 3,484 ± 142 | 2,678 ± 192 | 329 ± 52 | 0.016 ± 0.030 | 0,46 |
|  | Ac-20 | 6,846 ± 253 | 2,268 ± 2,079 | 135 ± 60 | 0.010 ± 0.008 | 0,75 |

As shown in Figure 3, cultures *C. acetobutylicum* and *C. beijerinckii* resulted in high lactic acid production. Table 4 highlights the production of volatile organic acids and the productivity of the most significant product, as well as solvent production and ethanol yield.

**Table 4.**
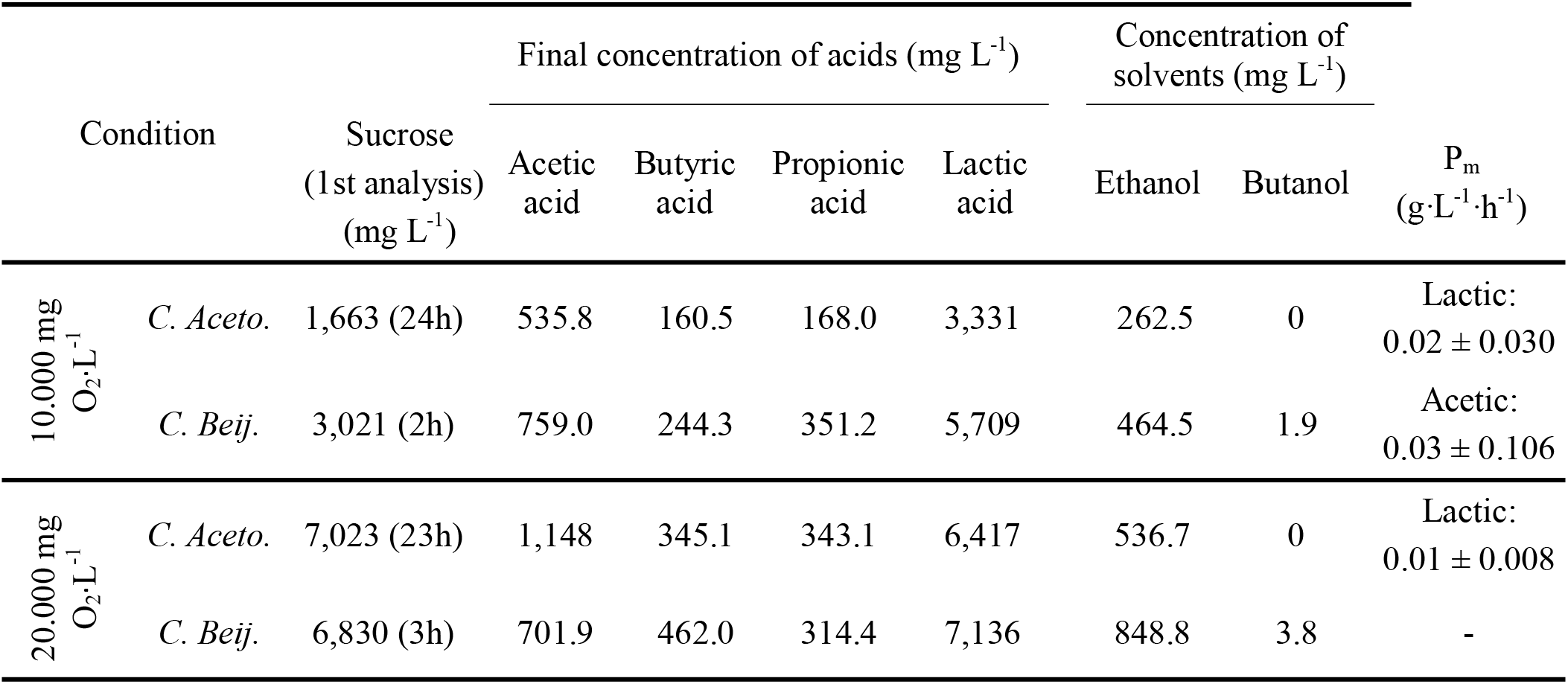
Initial concentrations of sucrose, final concentrations of volatile organic acids (acetic, butyric, propionic and lactic), final solvent concentrations (ethanol and butanol), average productivity (P_m_, g·L^-1^·h^-1^) and ethanol yield (Y_EtOH_).

As shown in Table 4, concentrations of COD of 10,000 mg O_2_·L^-1^ produced optimum lactic acid of 3,331 mg·L^-1^ and 5,709 mg·L^-1^ with respectively C. *acetobutylicum* and *C. beijerinckii.* Moreover, concentrations of COD of 20,000 mg O_2_·L^-1^ produced optimum lactic acid of 6,417 mg·L^-1^ and 7.136 mg·L^-1^ with respectively C. *acetobutylicum* and *C. beijerinckii*.

The complex substrate used allow to infer that both strains can adapt to the usage of vinasse as substrate to produce lactic acid. This use is presented as an alternative method of environmental waste management.

## 4 CONCLUSION

The study presented that both strains, *C. acetobutylicum* and *C. beijerinckii*, were able to produce lactic acid from a complex substrate without additional of vitamins, buffer solution and micronutrients. The production of lactic acid was favored presenting a final concentration of 7.136 g L-1, in the condition of 20,000 mg l-1 of COD. The results presented were highly influenced by pH and the acidogenic route was preponderant over the solventogenic route, despite the different inoculums and F/M-ratio studied.

## 5 ACKNOWLEGMENTS

The authors thank *Coordenação de Aperfeiçoamento de Pessoal de Nível Superior* (CAPES) for supporting this master study. Moreover, authors appreciate the collaboration of *Centro de Monitoramento e Pesquisa da Qualidade em Combustíveis, Biocombustíveis, Petróleo e Derivados* (CEMPEQC) for gas chromatography (CG) analysis focusing on butanol production.

